# Differential phoretic vector use among sympatric *Caenorhabditis* nematodes and an association with invasive nitidulid beetles in southwestern Germany

**DOI:** 10.64898/2025.12.16.694592

**Authors:** Ryan Greenway, Loel Dalan, Christian Braendle, Marie-Anne Félix, Siyu Serena Ding

**Affiliations:** Research Group Genes and Behavior, Max Planck Institute of Animal Behavior, 78464 Konstanz, Germany; Université Côte d’Azur, CNRS, Inserm, IBV, 06108 Nice, France; Institut de Biologie de l’École Normale Supérieure, Centre National de la Recherche Scientifique, 75005 Paris, France; Centre for the Advanced Study of Collective Behaviour, 78464 Konstanz, Germany

**Keywords:** Dauer, invasive species, interspecific interactions, coexistence, Coleoptera

## Abstract

Little is known about the natural history of many *Caenorhabditis* nematodes, despite their relationship to the model species *C. elegans*. While these nematodes rely on invertebrate vectors to disperse to new habitats (phoresy), vector use for most species has not been characterized. We surveyed the invertebrate community of a habitat containing three sympatric *Caenorhabditis* in southwestern Germany, determining differential and specific vector use for each of these co-occurring species. We documented a specific association between *C. apta* n. sp. and two species of invasive nitidulid beetles, and a particularly strong association of the nematode with the beetle *Stelidota geminata*, where we recovered more nematodes per individual beetle and a higher proportion of beetles carrying nematodes compared to the co-occurring *Epuraea ocularis*. Our results provide evidence for group dispersal using beetles in *C. apta*, supporting previous observations of collective dispersal behavior in this species, and establish a starting point for further dissecting the evolutionary and mechanistic causes and consequences of interactions between *Caenorhabditis* nematodes and their vector species in ecologically relevant conditions.

## Introduction

Free-living nematodes are some of the most abundant and diverse animals on Earth, and have reached their ubiquitous status despite their small size and limited modes of locomotion (Schratzberger *et al*. 2019). Many free-living nematodes achieve relatively wide distributions for their body size as the result of phoresy, hitchhiking to new habitats by temporarily attaching to a larger and more mobile interspecific vector (Bartlow & Agosta 2021). Phoresy in nematodes is often facilitated by an ability to enter diapaused or quiescent states, during which they can endure starvation and resist environmental extremes over extended time periods, allowing them to safely attach to vectors and disperse over greater distances than possible by crawling alone (Vlaar *et al*. 2021). The specificity of the relationship between phoretic nematodes and vector species varies widely across taxa, and is likely an important driver of nematode diversification and ecological success (Giblin-Davis *et al*. 2013; Bubrig & Fierst 2021; Vlaar *et al*. 2021).

*Caenorhabditis* nematodes are a key resource used to fuel a wide range of biological discovery over the last century, particularly the laboratory model organism *C. elegans* (Corsi *et al*. 2015; Meneely *et al*. 2019; Ambros *et al*. 2025). Despite their importance, the study of ecology and evolution in natural *Caenorhabditis* populations has only recently received significant attention thanks to the discovery of the natural habitat of these nematodes in the 2010s (Kiontke *et al*. 2011; Cutter 2015; Frézal & Félix 2015). We now know that in the wild *Caenorhabditis* are microbivorous nematodes with a boom-bust life cycle on ephemeral, microbe-rich resource patches, mostly decaying plant material (Kiontke *et al*. 2011; Cutter 2015). In the presence of stressors (*e.g.* starvation, high population density), *Caenorhabditis* nematodes develop a stress-resistant dauer larval form that facilitates dispersal and colonization of new patches after the rapid exhaustion of microbial resources (Barrière & Félix 2007; Félix & Duveau 2012; Schulenburg & Félix 2017; Vlaar *et al*. 2021). Experiments have suggested that *Caenorhabditis* appear to be reliant on phoretic associations with invertebrate carriers to colonize new resource patches (Li *et al*. 2014; Petersen *et al*. 2015; Sloat *et al*. 2022). Although phoretic dispersal is a key component of the life cycle and ecology of *Caenorhabditis*, interactions with potential dispersal vectors remain unknown for the vast majority of *Caenorhabditis* species.

A variety of invertebrate species have been documented carrying *Caenorhabditis* nematodes, with some species found in specialized associations with particular invertebrate taxa and others appearing to be opportunistic or generalists. The best characterized interaction between *Caenorhabditis* and vector is that of *C. japonica* with the shield bug *Parastrachia japonensis,* where the nematode is found exclusively in association with this insect throughout its life-cycle, and has not been found on other invertebrates despite examination of numerous co-occurring taxa (Yoshiga *et al*. 2013). Though not as well characterized as the relationship between *C. japonica* and its vector, *C. niphades* and *C. inopinata* also have specific relationships with particular insects (*Niphades variegatus* weevils and *Ceratosolen spp.* fig pollinating wasps, respectively; (Woodruff & Phillips 2018; Sun *et al*. 2022). Other *Caenorhabditis* may have similarly specific relationships with vector species or taxonomic groups, having been originally isolated and described in association with particular insects (Figure 1), such as *C. drosophilae* with the fruit fly *Drosophila nigrospiracula* (Kiontke 1997), *C. angaria* with the weevils *Metamasius hemipterus* and *Rhynchophorus palmarum* (Sudhaus *et al*. 2011), *C. plicata* with carrion beetles (Volk 1950)*, C. auriculariae* with fungus-feeding *Platydema spp.* beetles (Dayi *et al*. 2021), and *C. monodelphis* with the ciid beetle *Cis castaneus* (Slos *et al*. 2017). However, for these species there does not appear to have been systematic examination of other co-occurring organisms that could also act as vectors.

**Figure 1.**
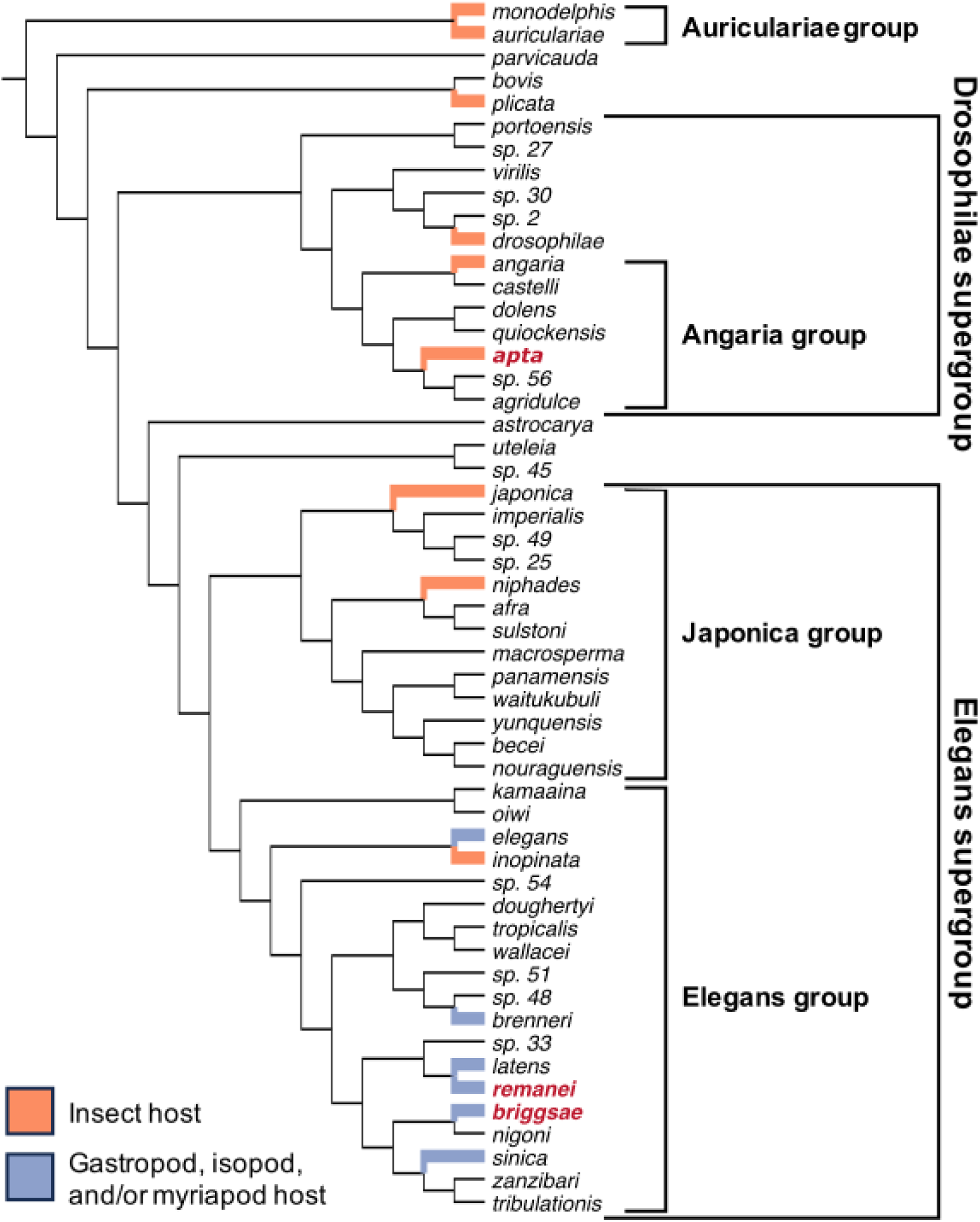
Phylogeny of *Caenorhabditis* nematodes, modified from Fusca *et al*. (2025), with the species groups and supergroups mentioned in this study indicated. Branches are colored by vector taxonomic group for species with documented associations, and the three species included in this study are indicated by bold red text.

In contrast to the specific interactions mentioned above, several *Caenorhabditis* species in the Elegans species group (Figure 1) have regularly been found in association with various species of terrestrial invertebrates, including *C. latens* with isopods (Dey *et al*. 2012)*, C brenneri* with gastropods (Diano *et al*. 2022)*, C. briggsae* (Barrière & Félix 2005; Félix & Duveau 2012; Félix *et al*. 2013; Petersen *et al*. 2015) and *C. sinica* (Wang *et al*. 2010) with isopods and gastropods, and *C. remanei* (Fitch *et al*. 1994; Baird 1999; Petersen *et al*. 2015) and *C. elegans* (Barrière & Félix 2005, 2007; Caswell-Chen *et al*. 2005; Félix & Duveau 2012; Ross *et al*. 2012; Petersen *et al*. 2015) with isopods, gastropods, and myriapods. Both *C. briggsae* and *C. elegans* have also been occasionally isolated from insects (Kiontke & Sudhaus 2006; Félix & Duveau 2012), and *C. elegans* dauers will attach to and disperse on various invertebrate groups, as well as inanimate materials, in laboratory experiments (Lee *et al*. 2011, 2017; Chiba *et al*. 2023; Williams *et al*. 2023; Perez *et al*. 2025). Consequently, it has been hypothesized that these species may be largely opportunistic in their use of phoretic vectors, exhibiting little specificity or preference for different taxonomic groups (Schulenburg & Félix 2017; Petersen *et al*. 2023). Under this scenario, it would be expected that opportunistic *Caenorhabditis* species could be recovered from many potential vectors frequently encountered in and around their habitats. Only one study has surveyed the broader invertebrate communities alongside *Caenorhabditis-*carrying gastropods, isopods, and myriapods but did not observe any additional invertebrate associations with *Caenorhabditis*, indicating there may be some degree of vector specificity for these nematodes (Petersen *et al*. 2015). Despite these described cases, vector use has not been characterized in any detail for the majority of the >80 known *Caenorhabditis* species.

In southwestern Germany, three species of *Caenorhabditis* can be found co-occurring at small spatial scales, even being found together on the same rotting fruits (Perez *et al*. 2025): *C. briggsae*, *C. remanei*, and *C.* sp. 8 described here as *C. apta* n. sp. (see Appendix 1), a little-studied species in the Angaria group of *Caenorhabditis* nematodes (Figure 1). Repeated sampling of the orchard meadows around the University of Konstanz campus has revealed that *C. apta* is the most abundant species in the area across multiple years; however, it remains unknown what vector(s) *C. apta* uses for dispersal. To determine whether these three *Caenorhabditis* species rely on the same or different vectors for dispersal, we conducted a broad survey of invertebrates found around rotting fruits. If any species were truly opportunistic in vector use, we would expect to recover them from a broad range of invertebrate groups surveyed from their habitat. Instead, we found that each *Caenorhabditis* species was isolated only from a subset of the sampled invertebrate taxa. Based on the results from this general survey, we conducted a more detailed survey of two locally abundant species of nitidulid beetles that served as vectors for *C. apta*. We also assessed the life stage, number of individuals, and sex ratio of *C. apta* on these two beetle species across multiple sites and across a three-month period at a single site to further characterize interactions between *C*. *apta* and its vectors.

## Materials and Methods

### General invertebrate collection and nematode extraction

To determine which invertebrate taxa serve as vectors for sympatric *C. briggsae, C. remanei*, and *C. apta*, we opportunistically collected invertebrates found on and around rotting fruits in orchard meadows near the University of Konstanz campus in Baden-Württemberg, Germany from June 20, 2025 to October 24, 2025 (Table 1). Gastropods and large arthropods were individually collected into Whirl-pak ® bags (The Artemis Corporation, USA) or 2 mL microcentrifuge tubes, while smaller arthropods (*e.g.* dipterans, coleopterans, hymenopterans) were collected using an aspirator and deposited into 2 mL microcentrifuge tubes. During this general survey, invertebrates were identified to order or higher taxonomic levels, with the exception of nitidulid beetles. Due to their frequent occurrence relative to other taxa, we identified nitidulid beetles to the species level based on keys for the beetle fauna of Europe (Lompe 2002; Benisch 2023). Invertebrates were brought back to the lab within two hours for processing. Invertebrates were individually added to either 6 or 9 cm petri dishes containing nematode growth media (NGM) seeded with a 200 µL spot of *E. coli* OP50. Petri dishes were sealed with Parafilm ® (Bemis Company, USA), left at 20° C on the benchtop in a climate-controlled laboratory, and monitored for 48 hours to determine if any nematodes emerged onto the bacterial lawn. Emerged nematodes were identified as *Caenorhabditis* based on morphology using a dissecting microscope (Stemi 508, Zeiss, Germany) fitted with 2.0x front optics. Following Barrière & Félix (2006), we identified *Caenorhabditis* based on the presence of both middle and basal pharyngeal bulbs, a clear grinder in the basal pharyngeal bulb, relatively light brown and homogeneously colored gut, a relatively short rectum, centrally located vulva (in females/hermaphrodites), a long and pointy tail in females/hermaphrodites, and a blunt tail with a fan-shaped bursa in males. We were also able to identify some additional nematode groups to the genus level, in particular *Pristionchus* based on the presence of well-developed teeth, the lack of a grinder in the terminal bulb of the pharynx, large and “dumpy” body shape, and clearly visible longitudinal striations in the cuticle (Barrière & Félix 2006; Ragsdale *et al*. 2015). *Sheraphelenchus* were distinguishable based on their diagnostic tail morphology, possessing a digitiform spike with a finely pointed tip, particularly evident in males (Hunt 1993; Kanzaki & Tanaka 2013). Lastly, a few individuals belonging to family Panagrolaimidae were able to be identified based on their long cylindrical pharynx, terminal pharyngeal bulb, and conical tail (Van Driessche *et al*. 2003). The identity of *Caenorhabditis* that emerged over the 48-hour period was determined (see below) and recorded for each sample.

**Table 1.**
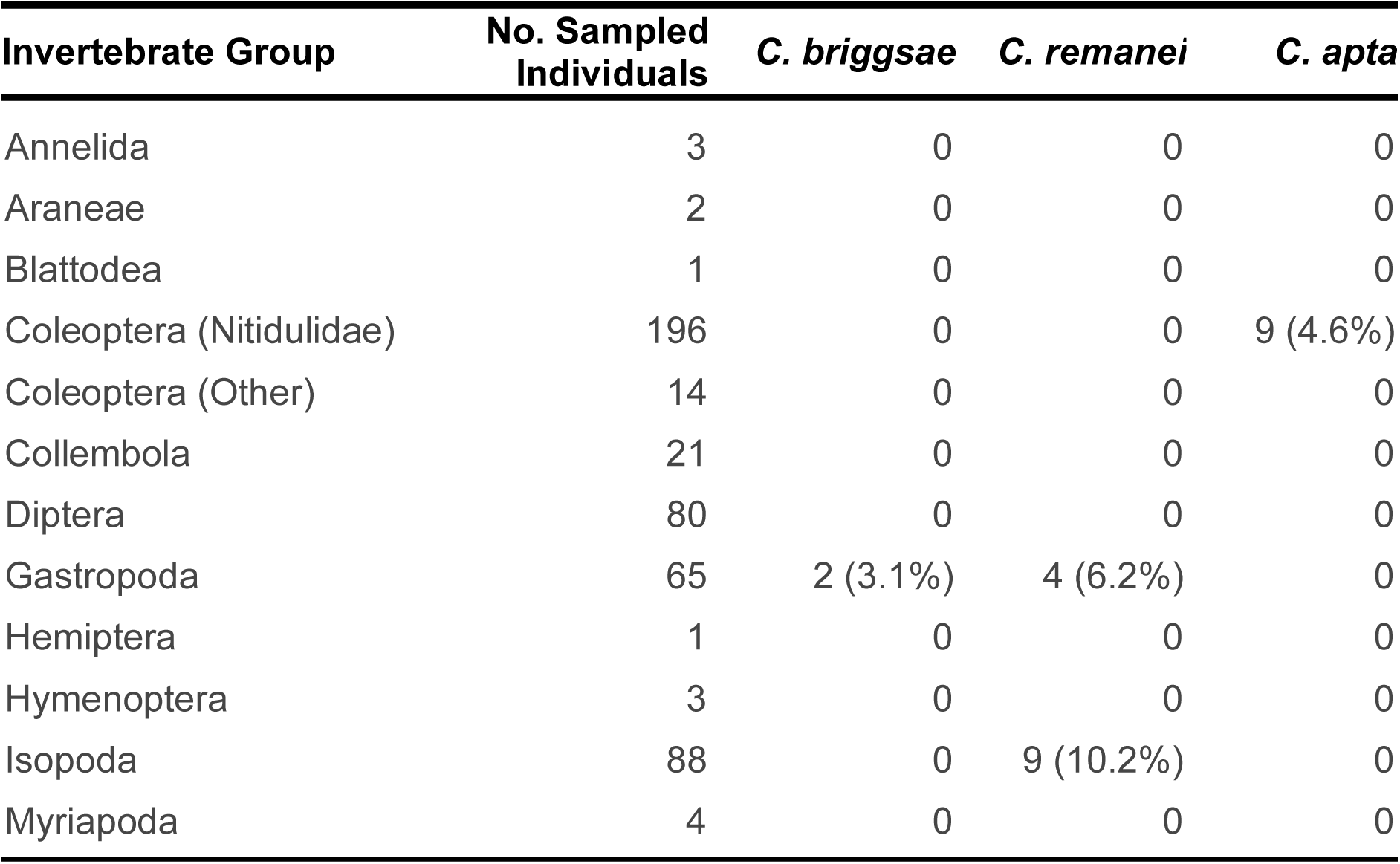
Summary of *Caenorhabditis* species isolated from different invertebrate groups around the University of Konstanz orchard meadows.

### Identification of Caenorhabditis species

We identified *Caenorhabditis* to the species level based on morphology, behavior, and reproductive mode using a Stemi 508 dissecting microscope (Zeiss, Germany) fitted with 2.0x front optics. Previous sampling of this area confirmed the presence of *C. remanei*, *C. briggsae*, and *C. apta* (Perez *et al*. 2025), which can be distinguished from each other based on established diagnostic criteria visible under a dissecting microscope, summarized in Appendix 2 (Barrière & Félix 2006; Kiontke *et al*. 2011; Sloat *et al*. 2022). After disembarking the invertebrates, nematodes were allowed to exit the dauer stage and resume development in order to facilitate identification. Upon reaching sexual maturity, nematodes were examined under the dissecting microscope and monitored for reproduction. We identified *C. briggsae* based on its self-fertilizing hermaphroditic reproduction and an absence of males, in contrast to the gonochoristic reproduction of the other two species. Of the remaining two species, we identified *C. apta* by its relatively short and “dumpy” body shape, spiral mating behavior, a narrow fan on the male tail, and stout, amber-colored male spicules. In contrast, *C. remanei* was identified based on its longer and slimmer body shape, parallel mating behavior, wide male tail fan, and less pronounced male spicules. In addition to these established criteria, we also observed that *C. remanei* tends to form large aggregations on bacterial lawns and readily burrow into the NGM compared to the other two species, while *C. briggsae* forage individually and tend to remain on the agar surface in the presence of food. Any uncertain species identifications were confirmed with mating tests, where isolated males or offspring generated from isolated mothers were individually transferred to a new 35 mm petri dish containing nematode NGM seeded with a 100 µL spot of *E. coli* OP50 and the opposite sex of known species identity to be assessed for reproduction over 72 hours, as each of these species is incapable of interbreeding with the other two (Greenway *unpublished data*, (Ting *et al*. 2014). In the case of putative hermaphrodites, virgin worms were plated individually and assessed for reproduction over 72 hours. Likewise, in cases where only a single female *Caenorhabditis* individual emerged from a given sample and did not reproduce 24 hours post-maturation, these individuals were transferred to new plates and provided with adult males of either *C. remanei* or *C. apta* (pairing based on body shape of the isolated individual) and monitored for reproduction over 72 hours to confirm species identity.

### Nitidulid beetle dissection and quantitative assessment of Caenorhabditis occupancy

Once we had identified beetles in the family Nitidulidae as the only invertebrates associated with *C. apta* (Table 1), we conducted more intense sampling of *Stelidota geminata* and *Epuraea ocularis*, two nitidulid species frequently observed and collected on rotting fruits in our prior sampling. To determine the species, life stage, number of individuals, sex ratio, and locations of nematodes on both beetle species, we performed dissections of adult *E. ocularis* and *S. geminata* collected from 6 locations in Baden-Württemberg, Germany (Table 2). Beetles were collected from rotting fruits using an aspirator, individually placed in 2 mL microcentrifuge tubes, and stored on ice for a minimum of one hour prior to sacrifice by removing the head from the thorax. The beetles were dissected under a stereomicroscope on a 6 cm petri dish containing NGM seeded with 200 µL *E. coli* OP50. We first examined the external surface of the beetle for the presence of nematodes before removing both elytra and transferring them together to a new NGM petri dish, leaving the rest of the corpse on the original dish. We then examined the internal surface of the elytra and the exposed abdomen separately, washing 20 µL of M9 buffer over each sample to loosen attached nematodes, at which point the life stage (larvae, dauer larvae, or adult) of visible nematodes was recorded. Petri dishes were sealed with Parafilm ® and left on the benchtop in a climate-controlled laboratory at 20° C. After 24 hours, the species, total number, and, for a subset of samples, the proportion of males present in the detached nematode population was separately recorded for the elytra and corpse of each beetle. The proportion of males on the beetle could not be determined in all cases, for example when some individuals remained as dauers for several days while others exited dauer rapidly and began to reproduce within 24 hours, resulting in mixed generation larval populations. We further monitored the corpse of each beetle for an additional 48 hours (72 hours total) to qualitatively determine the presence of additional nematode species that emerged following beetle death. To assess seasonal patterns of nematode occupancy and abundance on the beetles, we also collected each beetle species from a single location near the University of Konstanz (Egg Orchard; Figure 2A) two to three times per month from September 4, 2025 to November 10, 2025. Beetles were no longer found on or around fruits after the first freeze in mid-November.

**Figure 2.**
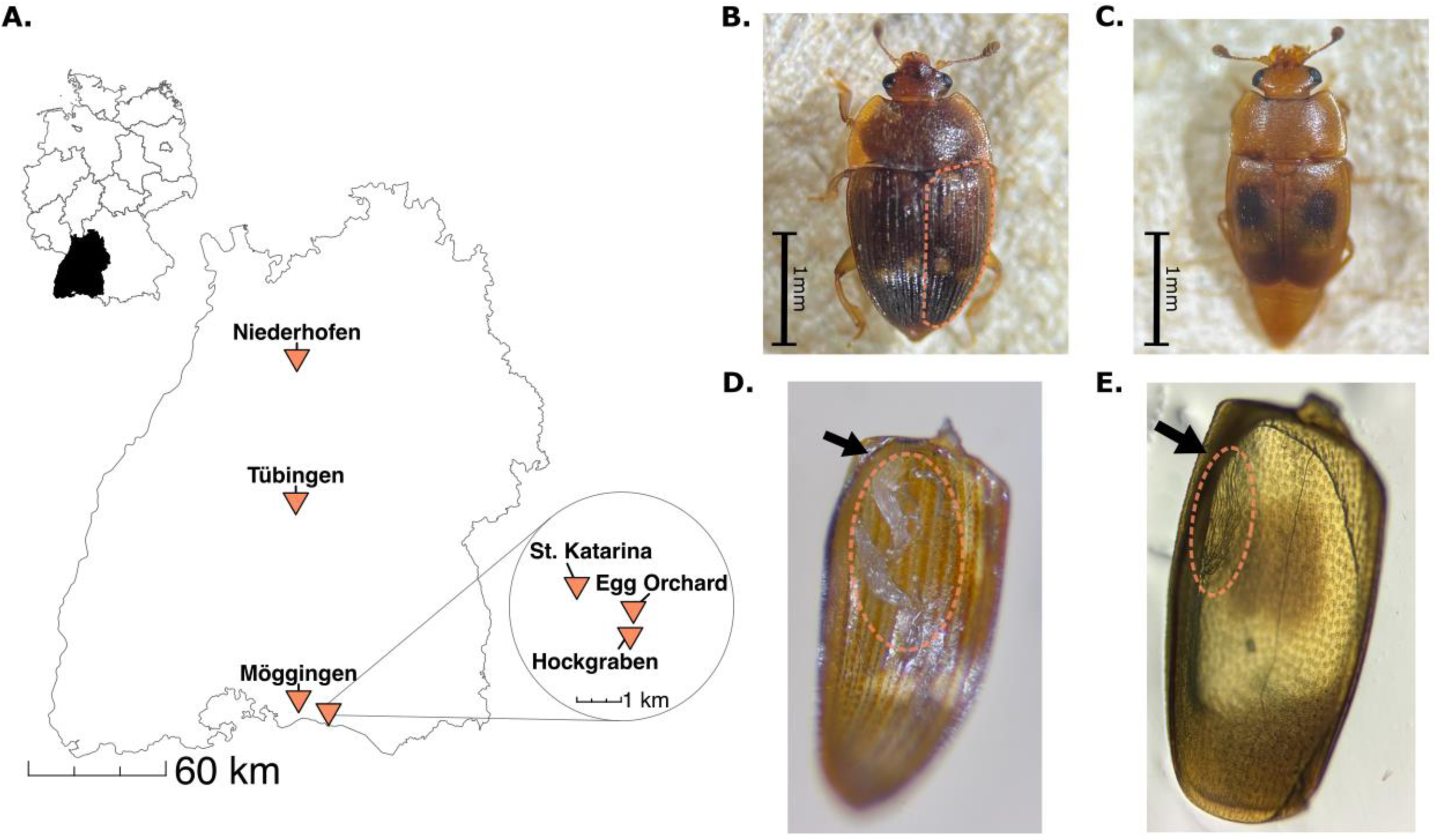
Collection sites and focal beetle species from this study. (**A**) *Stelidota geminata* and *Epuraea ocularis* were collected from six locations in Baden-Württemberg, Germany. The inset map depicts sites around the University of Konstanz campus. (**B**) *Stelidota geminata*, with the right elytron outlined. (**C**) *Epuraea ocularis*. (**D,E**) Clusters of dauer larvae (circled area) attached to the inner surface of the right elytron of *S. geminata* (>20 *C. apta,* **D**) or *E. ocularis* (>30 *Sheraphelenchus* sp., **E**). The epipleuron is indicated by an arrow in both images.

**Table 2.** Summary of among site variation in the number of beetles carrying *C. apta* dauers, the location of dauers on the beetles, and the sex ratio of *C. apta* from beetles.

| Site | Species | No. beetles | No. positive | Dauers/beetle<br>Mean ± SE | Dauers on elytra<br>Mean ± SE | Dauers on body<br>Mean ± SE | No. beetles<br>(Sex Ratio) | Sex Ratio<br>Mean ± SE |
| --- | --- | --- | --- | --- | --- | --- | --- | --- |
| Egg Orchard | *Epuraea ocularis* | 61 | 25 (41.0%) | 1.87 ± 0.53 | 1.84 ± 0.51 | 0.03 ± 0.02 | 13 | 0.34 ± 0.13 |
| | *Stelidota geminata* | 61 | 47 (77.0%) | 10.52 ± 3.37 | 9.62 ± 2.85 | 0.90 ± 0.58 | 30 | 0.47 ± 0.06 |
| Hockgraben | *Epuraea ocularis* | 30 | 10 (33.3%) | 1.37 ± 0.60 | 0.77 ± 0.23 | 0.60 ± 0.47 | 9 | 0.38 ± 0.14 |
| | *Stelidota geminata* | 30 | 24 (80.0%) | 10.83 ± 2.68 | 9.23 ± 2.45 | 1.60 ± 0.72 | 22 | 0.52 ± 0.05 |
| Möggingen | *Epuraea ocularis* | 35 | 8 (22.9%) | 0.51 ± 0.22 | 0.46 ± 0.21 | 0.06 ± 0.04 | 8 | 0.17 ± 0.09 |
| | *Stelidota geminata* | 35 | 18 (51.4%) | 2.34 ± 0.65 | 2.29 ± 0.65 | 0.06 ± 0.06 | 17 | 0.54 ± 0.08 |
| Niederhofen<sup>‡</sup> | *Epuraea ocularis* | 1 | 1 (100.0%) | 1.00 |  |  |  |  |
| | *Stelidota geminata* | 3 | 3 (100.0%) | 7.67 ± 5.24 | | | | |
| St. Katarina | *Epuraea ocularis* | 30 | 16 (53.3%) | 2.00 ± 0.59 | 1.70 ± 0.57 | 0.30 ± 0.09 | 12 | 0.49 ± 0.14 |
| | *Stelidota geminata* | 29 | 23 (79.3%) | 8.24 ± 1.91 | 7.59 ± 1.76 | 0.66 ± 0.38 | 13 | 0.48 ± 0.12 |
| Tübingen | *Epuraea ocularis* | 20 | 3 (15.0%) | 0.35 ± 0.22 | 0.35 ± 0.22 | 0 | 3 | 0.42 ± 0.30 |
| | *Stelidota geminata* | 39 | 20 (51.3%) | 4.56 ± 1.10 | 4.56 ± 1.10 | 0 | 20 | 0.54 ± 0.05 |
| All Sites<sup>‡</sup> | *Epuraea ocularis* | 177 | 63 (35.6%) | 1.36 ± 0.24 | 1.19 ± 0.22 | 0.18 ± 0.08 | 45 | 0.36 ± 0.06 |
| | *Stelidota geminata* | 197 | 135 (68.5%) | 7.56 ± 1.20 | 6.92 ± 1.05 | 0.64 ± 0.22 | 102 | 0.51 ± 0.03 |

<sup>‡</sup>Beetles from Niederhofen were excluded from analysis due to low sample sizes
SE = standard error

### Global distribution patterns of C. apta and its nitidulid vectors

Both nitidulid species we found carrying *C. apta* have invaded the European continent and spread rapidly since the 2000s (Konzelmann 2001; Callot 2007; Jelínek *et al*. 2016; Bibin 2017; Stan 2019; Grujić *et al*. 2020; Lasoń *et al*. 2023), leading us to ask whether this nematode may have been introduced to Europe alongside either of these species. To determine whether there is evidence that *C. apta* originated within the native range of either species, we examined the known distribution of *C. apta*, taking into account collection dates and geographic context. Specifically, we assessed whether *C. apta* had been collected in Europe, the Americas (the native range of *S. geminata*), or Southeast Asia and the western Pacific (the native range of *E. ocularis*). To this end, we compiled available sampling records from 1990–2024, including both published and new data, for *C. apta* (N = 55 strains) and other members of the Angaria clade (*C. quiockensis*, *C. castelli*, *C. angaria*, *C.* sp. 56, and *C. dolens*; N = 50 strains).

If *C. apta* was also recently introduced to Europe, we would expect to see collections of *C. apta* from the native range of one of these beetles, as well as detections in Europe starting in the mid to late 2000s, coinciding with the spread of the beetles. Alternatively, if the nematode is native to Europe, we would expect to see earlier detections of this species in Europe, prior to the spread of the beetles, but not necessarily overlapping distributions in the native range of either beetle species. Alternatively, *C. apta* may be a cosmopolitan species, in which case, we would expect to see a widespread distribution and no temporal pattern to this species detection in different geographic contexts. To provide a phylogeographic context to the distribution of *C. apta*, we also compared the known distributions of its closest relatives, species in the Angaria group. If many or all of the other species in the Angaria group are concentrated in a particular geographic area, this could indicate that *C. apta* also originates from the area. On the other hand, if the Angaria group species are cosmopolitan in distribution, it may indicate a similar natural range for *C. apta*. We also downloaded occurrence data for *S. geminata* and *E. ocularis* from GBIF (2025) to compare the geographic and temporal distributions of these two species with collection data for *C. apta*.

### Statistics

We analyzed all data in R v4.3.2 (R Core Team 2023). To test whether the proportion of individuals carrying *C. apta* differs across beetle species and sites, we used a generalized linear model (GLM) as implemented in the stats package (R Core Team 2023), with presence/absence of *C. apta* as a binomial response variable and a logit link function, including beetle species and site as factors. To determine whether the number of extracted *C. apta* individuals differed between the sampled body parts of the two beetle species across sites, we used a generalized linear mixed model (GLMM) assuming a negative binomial response variable (nematode counts) and log link function with the glmmTMB package (Brooks et al. 2017), using beetle species, body part, and site as factors, and the interaction between species and body part. We also included a random effect of the individual ID nested within site to account for non-independence among body parts from the same individual. To test if the sex ratio of *C. apta* individuals on beetles differed between species or sites, we modeled the proportion of male nematodes to total nematodes per beetle using a binomial GLM with a logit link function in the *stats* package, including beetle species and site as factors. The interaction between species and site was non-significant for the GLMs investigating differences in the number of beetles carrying *C. apta* and sex ratio differences, so were excluded from the final models. For these among site analyses, beetles collected from the Egg Orchard site were subsampled to better match the collection dates and sample sizes from other sites, including only individuals collected from September 15 to November 1, 2025. Additionally, the Niederhofen site was excluded from all analyses due to the low number of beetles sampled from this location.

We also used GLMs to test whether the number of individual beetles carrying *C. apta* and the number of *C. apta* individuals extracted from the two beetle species differed over time at the Egg Orchard site. We used a GLM modeling *C. apta* presence/absence on each beetle as a binomial response variable and a logit link function with the *stats* package, including beetle species as a factor and sampling date as a number (binning samples collected within 5 days of each other). We used a GLM with *C. apta* counts per beetle modeled as a negative binomial response with a log link function in the glmmTMB package, including beetle species and sampling date as factors. The non-significant interaction between species and date was excluded from both models.

## Results

### General invertebrate survey for Caenorhabditis

We first surveyed a broad sample of invertebrates found on and around rotting fruits near the University of Konstanz, noting the presence or absence of the three *Caenorhabditis* species on each individual to determine which taxa serve as vectors for *Caenorhabditis* nematodes in southern Baden-Württemberg. In total, 478 individual invertebrates from various taxonomic groups were included in this general survey (Table 1). Confirming the results of previous studies (Fitch *et al*. 1994; Baird 1999; Dey *et al*. 2012; Félix & Duveau 2012; Félix *et al*. 2013; Petersen *et al*. 2015), *C. briggsae* and *C. remanei* were both found on gastropods, and *C. remanei* on isopods. *C. briggsae* was isolated from two slugs (3.1% of gastropods), while *C. remanei* was isolated from four slugs (6.2% of gastropods) and 9 isopods (10.2% of isopods). The two species were never found occurring on the same vector individual in this survey. In contrast, *C. apta* was never found on gastropods or isopods, but instead was only found in association with beetles in the family Nitidulidae, being isolated from 9 nitidulid beetles (4.6% of all nitidulids) in the general survey. This, however, likely represents an underestimation of the total occupancy rate of *C. apta* on nitidulids based on closer inspection of these beetles (see detailed survey below) and laboratory observations. We found *C. apta* occurring in several independent nitidulid breeding cultures started from beetles that were deemed “nematode free” after initial screening on agar plates. Several days after placing these beetles on sterilized apples, many of the cultures were covered in *C. apta* nematodes, indicating that *C. apta* may be reluctant to disembark vectors when presented only with *E. coli* OP50 as a food source. In light of this, we conducted closer inspection of isopods and gastropods to confirm that these taxa did not also carry *C. apta* or other *Caenorhabditis* that were reluctant to depart onto the provided media. A subset of the isopods and gastropods from the general survey (*n* = 34 and 15, respectively) were killed with a scalpel and washed in water, plated on NGM petri dishes with the washing liquid, and observed for approximately one week. We did not find any additional *Caenorhabditis* emerging from these individuals, confirming the absence of *C. apta* on these taxa. Additionally, *C. briggsae* and *C. remanei* were never found on nitidulid beetles, and no *Caenorhabditis* species were found from any other invertebrate group in the survey (Table 1).

### Quantitative assessment of C. apta from nitidulid beetles

As an association between *Caenorhabditis* and nitidulids has not been previously described, we further studied this relationship by conducting additional sampling of *E. ocularis* and *S. geminata* (Figure 2B and C), the two nitidulid beetles most often encountered in our study area, for quantitative characterization of *C. apta* from these beetles. We dissected an additional 177 *E. ocularis* and 197 *S. geminata* collected between September 15 and November 1, 2025 across six sites in Baden-Württemberg (Figure 2A; Table 2) to determine the lifestage, prevalence, and sex ratio of *C. apta* on individual beetles, as well as where on the beetles the nematodes could be found. We found that the abundance of *C. apta* significantly differed among body parts, beetle species, and sites. Inspection of both the elytra and remainder of the beetle corpse revealed that only dauer larvae were found on the beetles (Figure 2D). The number of *C. apta* observed was higher on the elytra than on the rest of the beetle body (β = 2.31 ± 0.33, z = 7.10, p < 0.001), with *S. geminata* having significantly higher abundances than *E. ocularis* on both the elytra and body (interaction β = 0.97 ± 0.40, z = 2.40, p = 0.016; species β = 1.14 ± 0.37, z = 3.07, p = 0.002; Table 2). Dauers were mostly found on the anterior portion of the elytra, stuck together into clusters in or near a cavity created by the epipleura (the portion of the elytron that folds back towards the abdomen (Goczał & Beutel 2023); Figure 2D and E). Attached dauers were largely motionless until prodded with a wire pick or wetted with M9 solution, at which point they began to wriggle and detach from each other. Despite concentration on the elytra, low numbers of *C. apta* dauers were also found having emerged from the rest of the beetle body in some cases (Table 2). The number of *C. apta* observed on beetles was significantly lower at Möggingen (β = −1.22 ± 0.34, z = −3.55, p < 0.001) and Tübingen (β = −0.99 ± 0.35, z = −2.84, p = 0.004) compared to the Egg Orchard site (Table 2).

We also examined the sex ratio of *C. apta* individuals emerged from each beetle (combined across body parts), finding that the proportion of males on each beetle was significantly lower from an expected 1:1 ratio (Huang *et al*. 2023) for *E. ocularis* (Intercept β = −0.68 ± 0.23, z = −2.93, p = 0.003) and was significantly different between species, closer to an even 1:1 for *S. geminata* (β = 0.53 ± 0.22, z = 2.46, p = 0.014; Table 2), but did not differ among sites (all z ≤ 1.59, all p > 0.110). Differences in the sex ratio of attached nematodes between the two species are likely driven by the low number of nematodes recovered per beetle for *E. ocularis* (Table 2).

In addition to carrying significantly more dauers per beetle, a higher proportion of *S. geminata* individuals were carrying *C. apta* dauers than were *E. ocularis* (β = 1.57 ± 0.24, z = 6.59, p < 0.001), with *C. apta* found on 68.5% of *S. geminata* individuals compared to 35.6% of *E. ocularis* across all sites (Table 2). However, there were also differences in the proportion of all beetle individuals carrying *C. apta* dauers among sites, with lower overall rates of nematode recovery in both Möggingen (β = −1.03 ± 0.33, z = −3.09, p = 0.002) and Tübingen (β = −1.20 ± 0.35, z = −3.40, p < 0.001) compared to the Egg Orchard site (Table 2). As these sites were only sampled a single time later in the season, it is impossible to distinguish if the lower dauer abundance and proportion of beetle vectors are the result of site-specific or temporal variation in this interspecific interaction.

### Temporal variation in nematode-beetle interactions

We examined temporal variation in the number of beetles carrying *C. apta* and the number of individual nematodes per beetle at a single site (Egg Orchard, Konstanz; 105 *E. ocularis*, 123 *S. geminata*) from September to November 2025 (Figure 3), spanning the season when *C. apta* is abundant and active in late summer until its disappearance in mid-autumn. Dauers of *C. apta* were found on a significantly higher number of *S. geminata* than *E. ocularis* over the entire period (β = 1.92 ± 0.31, z =6.25, p < 0.001), though the overall proportion of nitidulids carrying *C. apta* decreased over time (β = −0.43 ± 0.16, z =-2.73, p = 0.006). Likewise, *S. geminata* carried significantly more *C. apta* dauers over the whole season than did *E. ocularis* (β = 2.13 ± 0.24, z =8.97, p < 0.001), and the number of dauers on both species decreased over time (β = −0.50 ± 0.14, z =-3.71, p < 0.001). Though this survey misses early season and over-winter dynamics, we recover a consistent pattern of higher *C. apta* dauer occupancy on *S. geminata* than on *E. ocularis* across time, and expect a qualitatively similar pattern exists across the year, particularly since nitidulid beetles overwinter as adults.

**Figure 3.**
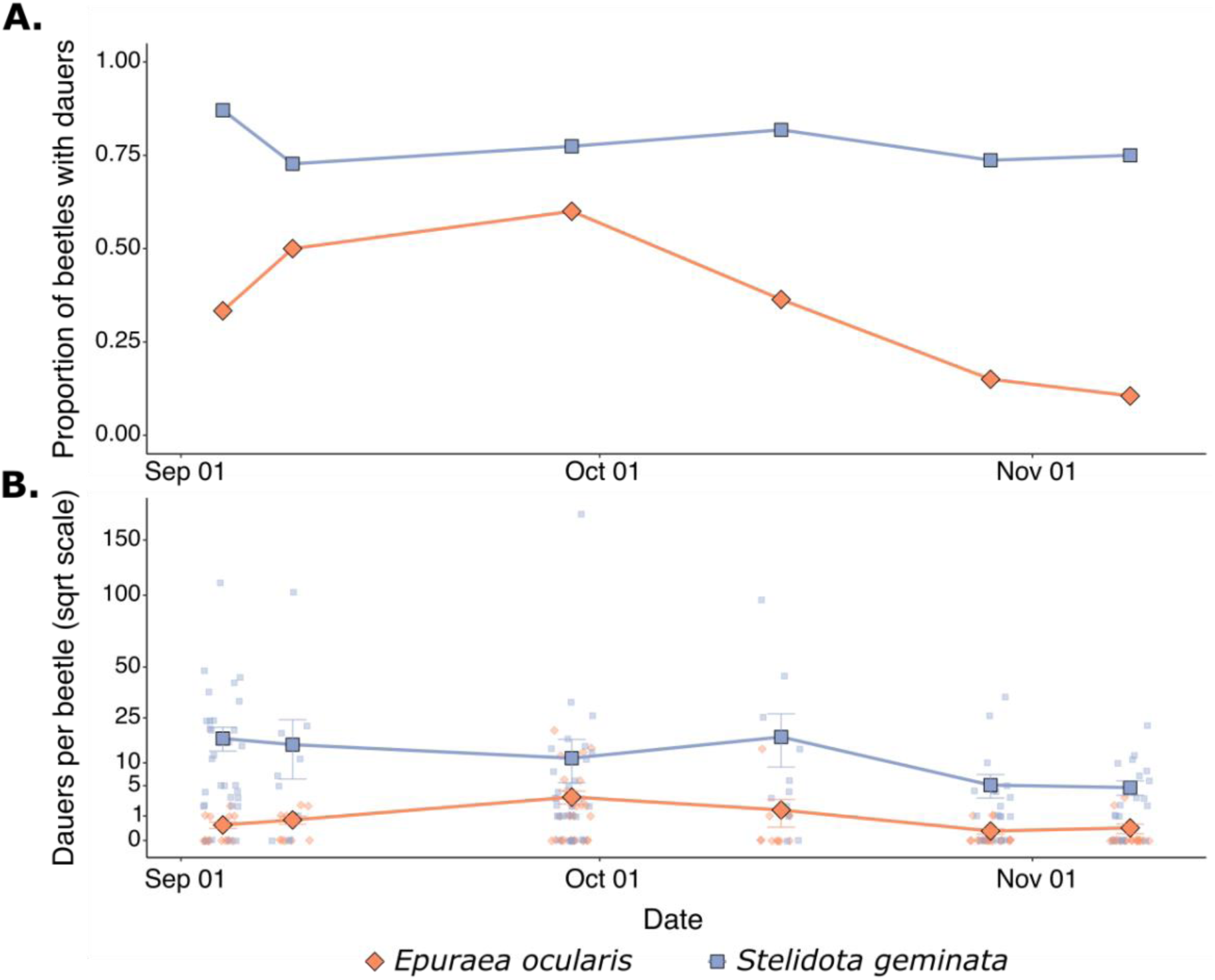
Temporal variation in (**A**) the proportion of beetles carrying *C. apta* and (**B**) the number of *C. apta* dauer larvae extracted per beetle (square root scaled) for two nitidulid beetle species. In **B**, large points represent the mean number of nematodes per beetle for each species with error bars depicting the standard error, small points represent the number of nematodes extracted from individual beetles.

### Other nematodes isolated from nitidulid beetles

While *C. apta* was the nematode species most often observed in our survey of these two nitidulid species, several additional nematode species were encountered on occasion. The next most frequently recovered was *Sheraphelenchus* (likely *S. sucus* based on frequent occurrence on fruits in the area [(Perez *et al*. 2025)]), a mycetophagous nematode in the family Aphelenchoididae (Kanzaki & Tanaka 2013). *Sheraphelenchus* were found on 62 *E. ocularis* (28.1% of sampled beetles) and 14 *S. geminata* (5.4% of sampled beetles), co-occurring with *C. apta* on 15 and 9 individual beetles, respectively. As seen with *C. apta*, *Sheraphelenchus* were also frequently observed clustered into large groups on the underside of the elytra of *E. ocularis* (Figure 2E). In addition to these commonly observed species, we also recovered a hermaphroditic *Pristionchus* species from 1 *E. ocularis* (0.45% of sampled beetles) and 2 *S. geminata* (0.77% of sampled beetles), and a hermaphroditic Panagrolaimid from 2 *E. ocularis* individuals (0.90% of sampled beetles), all co-occurring on beetles with *C. apta* with the exception of one of the Panagrolaimids.

### Global distribution patterns of C. apta and its nitidulid vectors

Beginning in 2007, *C. apta* was first isolated from sites across North America. In 2011, 1. *C. apta* was recorded for the first time from European collections. With the exception of *C. apta,* all other known species in the Angaria group of *Caenorhabditis* nematodes have only been collected from the Americas (Kiontke *et al*. 2011; Sudhaus *et al*. 2011; Ferrari *et al*. 2017; Stevens *et al*. 2019; Sloat *et al*. 2022). North America is also the native of range of *S. geminata* (Weber & Connell 1975), where it overlaps with the few available collections of *C. apta* (Figure 4). *E. ocularis*, on the other hand, is native to Southeast Asia and the western Pacific (Cline & Audisio 2011; Rittner & Nir 2013), where *C. apta* has never been detected. Furthermore, *C. apta* has a wide and overlapping distribution with the invasive range of *S. geminata* in Europe (Figure 4).

**Figure 4.**
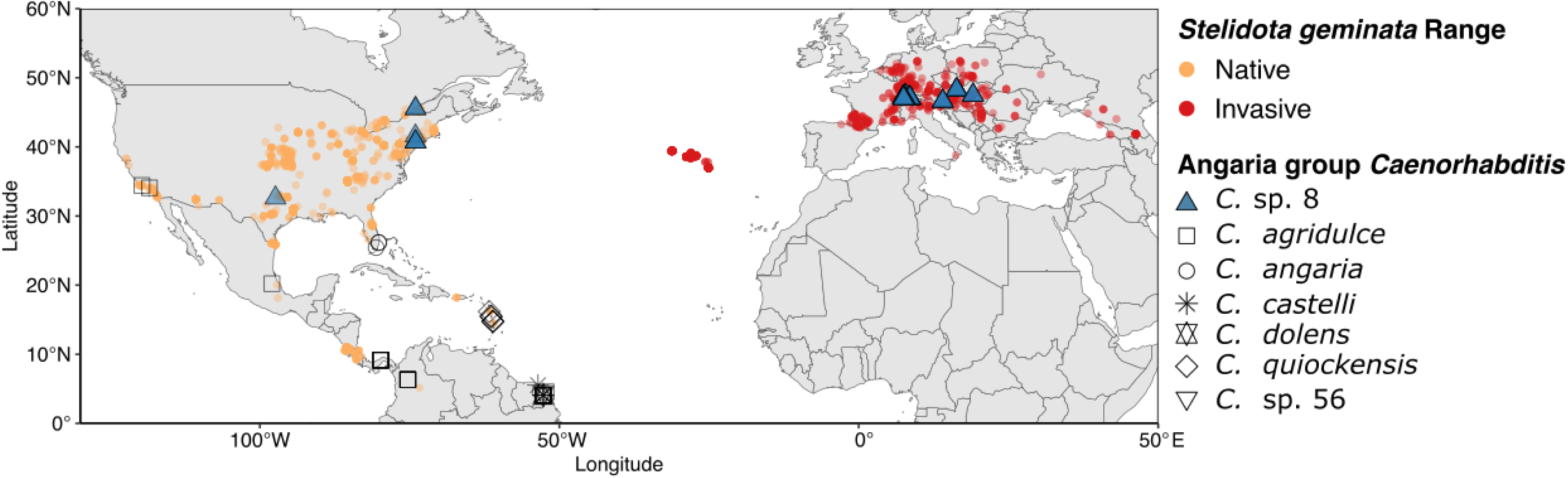
Global distribution of *S. geminata* and known occurrences of *Caenorhabditis* species in the Angaria group, including *C. apta* (blue triangles). *S. geminata* has spread from its native range in the Americas to Europe since the 2000s (orange versus red points; more intense coloration corresponds to multiple overlapping occurrences). As most Angaria group species have been described and sampled from French Guiana, data points heavily overlap in this region.

## Discussion

A better understanding of the relationships between *Caenorhabditis* nematodes and their phoretic vectors is necessary for characterizing the factors shaping the ecology and evolution of these model organisms. However, studies of variation in vector use among sympatric *Caenorhabditis* species, as well as basic characterization of vector use for most *Caenorhabditis* species, have remained limited. In this study, we quantified associations between three sympatric *Caenorhabditis* species and a variety of invertebrate taxonomic groups, finding variation in vector use among species, as well as a novel vector association for *C. apta*.

### Variation in vector use among sympatric Caenorhabditis species

Despite all three *Caenorhabditis* species co-occurring at small spatial scales, even on the same rotting fruits (Perez *et al*. 2025), we found variation in vector use among species. In our general survey of 478 invertebrates from the orchard meadows near the University of Konstanz, we found vector associations between *C. briggsae* and *C. remanei* with gastropods, as well as *C. remanei* with isopods, but none of the other surveyed taxa, consistent with vector associations previously reported for these species (Fitch *et al*. 1994; Baird 1999; Félix & Duveau 2012; Petersen *et al*. 2015). *Caenorhabditis apta* was never recovered from gastropods or isopods, instead only being found in association with nitidulid beetles, and particularly with *S. geminata*. It cannot be entirely ruled out that other beetle or insect species could act as vectors for *C*. *apta*, particularly given the low sample size for some taxonomic groups; however, based on the occurrence rates of the sampled invertebrates on and around rotting fruit in our study area, we are confident that nitidulid beetles are frequently used vectors in this area. The high occupancy of *C. apta* on nitidulids that we recorded across multiple sites in Baden-Württemberg may indicate a higher propensity or ability for dispersal in this species compared with the other two, perhaps mediated by underlying differences in dispersal-related behaviors as has been observed among *C. elegans* isolates (Lee *et al*. 2017). In line with this, the occurrence rates of *C. briggsae* and *C. remanei* on isopods and gastropods documented in our survey roughly corresponds with the results of a previous survey of vector-nematode associations from northern Germany (Petersen *et al*. 2015), potentially indicating species-specific dispersal strategies. Further work is necessary to determine the relationship between vector use, dispersal ability, and community dynamics.

Our findings also provide evidence that *Caenorhabditis*-invertebrate interactions are governed by mechanisms facilitating some degree of vector specificity. If Elegans group nematodes were entirely opportunistic in their vector use, we would expect to occasionally recover either species from the extremely common nitidulid beetles found on most fruits in the study area. However, neither *C*. *briggsae* nor *C. remanei* was found on any of the >450 nitidulid beetles sampled in this study. Likewise, *C. apta* was not recovered from any gastropods or isopods, despite frequent occurrence of these invertebrates on and around fruit containing *C. apta* dauers. Laboratory experiments dissecting vector preference for different invertebrate species among these *Caenorhabditis* could provide additional insight into the true degree of vector specificity for each species and the mechanisms governing how they are able to differentiate among co-occurring vectors. Preference for vector species has been experimentally demonstrated for the highly specialized *C. japonica*, mediated by chemotactic attraction to vector cues (Okumura *et al*. 2013; Okumura & Yoshiga 2014), while experiments with *C. elegans* have failed to find any evidence of attraction to the chemical profiles of known vector species (Archer *et al*. 2020; Petersen *et al*. 2023). Determining how chemosensory or other cues aid in vector detection and response will be of great value for better understanding the ecology, genetics, and evolution of *Caenorhabditis* nematodes.

### C. apta dispersal on nitidulid beetles

Detailed analysis of the association between *C. apta* and the two nitidulid beetles in our study area highlighted that *C. apta* dauers frequently disperse in mixed sex groups on the underside of beetles’ elytra. Several beetle-dispersed nematodes form similar clumps or clusters of individual worms on the inner surfaces of elytra (Penas *et al*. 2006; Shimizu *et al*. 2013; Kanzaki *et al*. 2019; Polyanina *et al*. 2019; Goczał & Beutel 2023). It remains unclear whether these elytral clusters are the direct result of group dispersal behaviors or form once multiple nematodes have individually attached to the vector. We have previously observed *C. apta* forming dauer towers in nature, as well as towers reacting to and briefly attaching to fruit flies in the laboratory (Perez *et al*. 2025); however, it remains to be tested how dauer towers directly interact with nitidulids, the apparent vectors of this species. Given the coordinated locomotion of dauers in the tower structure on rotten fruits, it is plausible that the elytra clumps we observe here are the direct results of towers, having collectively migrated underneath the elytra once attached to the beetle. However, our recovery of both individual dauers and groups of dauers from each beetle species indicates *C. apta* is can disperse on nitidulid beetles both as individuals and in groups, providing an excellent system for disentangling the benefits and costs of collective versus individual dispersal.

In addition to dispersing with members of the same species, we also observed co-occurrence of more than one nematode species on the same beetle. As the interactions between *C. apta* and co-occurring nematodes are yet unknown, it is unclear what the causes or consequences of multi-species phoresy may be. *Sheraphelenchus* and *Caenorhabditis* likely are not competing for resources, either feeding on fungal mycelia or free-living microbes, respectively. On the other hand, *Pristionchus* nematodes can be predatory, feeding on the larvae of other nematode species (Hiramatsu & Lightfoot 2023; Sommer 2025), and may take advantage of dispersing with other species in order to secure a meal upon arrival to a new habitat patch. Testing whether there are benefits to dispersing in mixed-species groups versus alone for any of these species would help shed light on whether co-occurrence is simply due to chance or if one species may seek out or avoid beetles carrying the others (Gupta & Borges 2021).

It is an open question whether either of these invasive nitidulid species is the natural vector of *C. apta*, with which it has expanded its range to Europe by hitchhiking together, or instead both of these beetle species are novel vectors *C. apta* has recently begun to use. Given the known distribution of all other Angaria group species, the co-distribution of *C.* apta and *S. geminata,* and the stronger association we detected between *C. apta* and *S. geminata* compared to *E. ocularis*, it is possible that *C. apta* is in fact a recent colonizer of Europe as a result of the expansion of its natural beetle vector from North America. Co-invasion on insect vectors has been documented for other phoretic nematodes (D’Anna & Sommer 2011), sometimes with severe ecological and economic consequences (Giblin-Davis *et al*. 2013). However, it is also possible that, while initially undetected in Europe due to a patchy distribution or the lack of targeted sampling, *C. apta* is indeed a native species that has recently been able to increase in abundance or locally expand its range following a switch to the use of invasive vector species, as has been observed in *Pristionchus* (D’Anna & Sommer 2011). Currently, the poor sampling of *C. apta* in North America, as well as a lack of ecological information for this species, hinders further study of the history of these interactions. Additional support for the invasion hypothesis will require documentation of a similar phoretic interaction in the native range of *S. geminata*, as well as population genetic analyses of *C. apta* from populations sampled from across the known range. If *C. apta* is an invasive species in Europe, this system would provide an excellent opportunity for studying the ecological and evolutionary consequences of range expansion in *Caenorhabditis* nematodes, such as how native *Caenorhabditis* species respond to a novel competitor.

## Conclusion

Invertebrate vector associations for *Caenorhabditis* nematodes in southwestern Germany correspond with current understanding of vector use in the genus. *C. briggsae* and *C. remanei* associate with terrestrial invertebrates, though the mechanisms governing this association remain unknown. The sympatric *C. apta*, however, was exclusively recovered from nitidulid beetles, exhibiting a strong association with *S. geminata* across both time and space. Further characterization of the association between *C. apta* and these invasive nitidulid species in Europe and the native ranges of the beetles will provide a better understanding of the ecology and evolution of this little-studied nematode species.

## Data Accessibility Statement

All data and code necessary to replicate the analysis and figures presented in this manuscript are available at https://github.com/SerenaDingLab/Greenway_et_al_Caen-Invert-Intxns.

## Competing Interests Statement

The authors have no competing interests to declare.

## Author Contributions

Conceptualization, data curation, formal analysis, visualization, and writing - original draft preparation: R.G. Investigation: R.G., L.D., C.B., and M.A.F. Writing - review and editing: R.G., L.D., C.B., M.A.F., and S.S.D. Supervision, project administration, resources, and funding acquisition: S.S.D.

## Acknowledgements

We would like to thank Anna-Lena Bucher, Narcís Font-Massot, Matthias Hermann, Berke Güner, and Christian Weiler for their help accessing and traveling to field sites. We thank all members of the Ding Lab at the MPI of Animal Behavior for constructive discussions of the results and feedback on the manuscript draft. This study was funded through the Max Planck Institute of Animal Behavior.

## Appendix 1. Description of *Caenorhabditis apta* sp. n. (Nematoda: Rhabditidae)

### Rationale for raising a new species

Most closely related *Caenorhabditis* species are highly similar morphologically; therefore, following Félix *et al*. (2014) we use the amenability of *Caenorhabditis* species to culturing and test crosses to delineate species based on the biological species concept. Following Félix *et al*. (2014), crosses were prioritized using genetic proximity. Building a consensus tree based on sequence alignments of 2,955 genes, Rockman *et al*. (2025) found the closest described species of *C. apta* QX1182 to be *C. agridulce* (Sloat *et al*. 2022). Sloat *et al*. (2022) reported that crosses between *C. apta* QX1182 males and *C. agridulce* QG555 females produced arrested embryos and a small number of L1 larvae that died, while the reciprocal cross only produced arrested embryos. *Caenorhabditis apta* thus defines a new biological species.

### Methods

We used scanning electron microscopy for documenting and comparing morphological features of *C. apta* with closely related species. QX1182 nematodes on NGM agar culture plates were resuspended and washed twice in M9 solution, and fixed overnight at 4°C in glutaraldehyde 2.5%, in M9 or 50 mM phosphate pH 7.0 depending on the batch. The fixed animals were rinsed twice in M9 and dehydrated through an ethanol series, pelleting them at each step at 1 g in a tube. The samples were processed through critical point drying and coating with 20 nm of Au/Pd, and observed with a JEOL 6700F microscope at the Ultrastructural Microscopy Platform of the Pasteur Institute.

### Species declaration

The electronic edition of this article conforms to the requirements of the amended International Code of Zoological Nomenclature, and hence the new names contained herein are available under that Code from the electronic edition of this article. This published work and the nomenclatural acts it contains have been registered in ZooBank, the online registration system for the ICZN. The ZooBank LSIDs (Life Science Identifiers) can be resolved and the associated information viewed through any standard web browser by appending the LSID to the prefix ‘‘http://zoobank.org/’’. The LSID for this publication is: LSID urn:lsid:zoobank.org:pub:E5238355-9049-4697- 9422-97C9F2EA5616. The electronic edition of this work was published in a journal with an ISSN. We provide pictures in Appendix Figs S1 and S2 to conform with the ICZN guidelines.

*Caenorhabditis apta* sp. n. Greenway, Dalan, Braendle, Félix, & Ding Zoobank Identifier

urn:lsid:zoobank.org:act:E679D6EE-FDB9-476C-A274-6712AF93AA26 = *Caenorhabditis sp. 8* in (Kiontke *et al*. 2011; Félix *et al*. 2014; Vielle *et al*. 2016; Sloat *et al*. 2022; Rockman *et al*. 2025)

The type isolate by present designation is QX1182, deposited at the *Caenorhabditis* Genetics Center (https://cgc.umn.edu/). The species is delineated and diagnosed by the fertile cross with the type isolate QX1182 in both cross directions, yielding highly fertile hybrid females and males that are interfertile and cross-fertile with their parent strains. The species reproduces through males and females. The ribosomal DNA sequence of *C. apta* QX1182 including ITS2 is accessible at NCBI with accession number JN636143 (Kiontke *et al*. 2011). This species differs by SSU, LSU and ITS2 DNA sequences (Kiontke *et al*. 2011) from all species listed in Tables 1 and 2 of (Félix *et al*. 2014; Huang *et al*. 2014; Ferrari *et al*. 2017; Slos *et al*. 2017; Kanzaki *et al*. 2018; Crombie *et al*. 2019; Stevens *et al*. 2019, 2020; Dayi *et al*. 2021; Sloat *et al*. 2022; Devi *et al*. 2025; Rockman *et al*. 2025). Note that these ribosomal DNA sequences may vary within the species (Kiontke *et al*. 2011). The type isolate was collected by Matthew Rockman from rotting tomatoes in New Jersey, USA, in July 2007 (Kiontke *et al*. 2011). Other isolates were found in decomposing fruits in USA and Germany (Kiontke *et al*. 2011; Perez *et al*. 2025), as well as on decomposing fruits in Switzerland, Austria, Czech Republic and Hungary and on nitidulid beetles in Germany in the present study. The mouth is endowed with the usual set of sensory organs disposed in a concentric manner, namely six labial sensillae, two amphids and six male-specific cephalic sensillae (Fig. A1A-C). The radial symmetry of the labial sensillae appears broken by the more external position of the two sensillae closest to the amphids (Fig. A1A-B). An inner flap is present on each of the six lips, as in *C. angaria* (Sudhaus *et al*. 2011) and *C. agridulce* (Sloat *et al*. 2022) (Fig. A1A-C). The male tail displays an anteriorly open and narrow fan with a smooth edge (Fig. A2B-D). The dorsal rays are in antero-posterior positions 4 and 7 (Fig. A2A-D). Papilliform phasmids are visible (Kiontke *et al*. 2011) (Fig. A2A). The spicule tips are thick and curved internally (Fig. A2E-F). The pre-cloacal sensillum is semi-circular and appears bordered anteriorly by an additional cuticular lip (Fig. A2G-H). In addition, a drawing of the male tail can be found in Kiontke *et al*. 2011. A Nomarski microscopy picture of the buccal cavity and a drawing of a spicule can be found in (Sloat *et al*. 2022). The males mate in a spiral position. The species is named after their ability to be connected and attached together in masses.

## Appendix 2.

**Table A1.** Summary of characters used to distinguish among sympatric.

| Character | *C. apta* | *C. briggsae* | *C. remanei* |
| --- | --- | --- | --- |
| *Body shape* | Short, dumpy | Long, slender | Long, slender |
| *Reproductive mode* | Gonochoristic | Hermaphroditic and selfing | Gonochoristic |
| *Mating behavior* | Spiral | - | Parallel |
| *Male morphology* | Stout, amber colored spicules<br>Narrow bursal fan | - | Smaller spicules<br>Wide bursal fan |
| *Foraging behavior* | Forms small aggregations on food<br>No observed burrowing on food | Solitary on food<br>No observed burrowing on food | Forms large aggregations on food<br>Burrows into NGM when food present |

---

**Appendix Figure A1.**
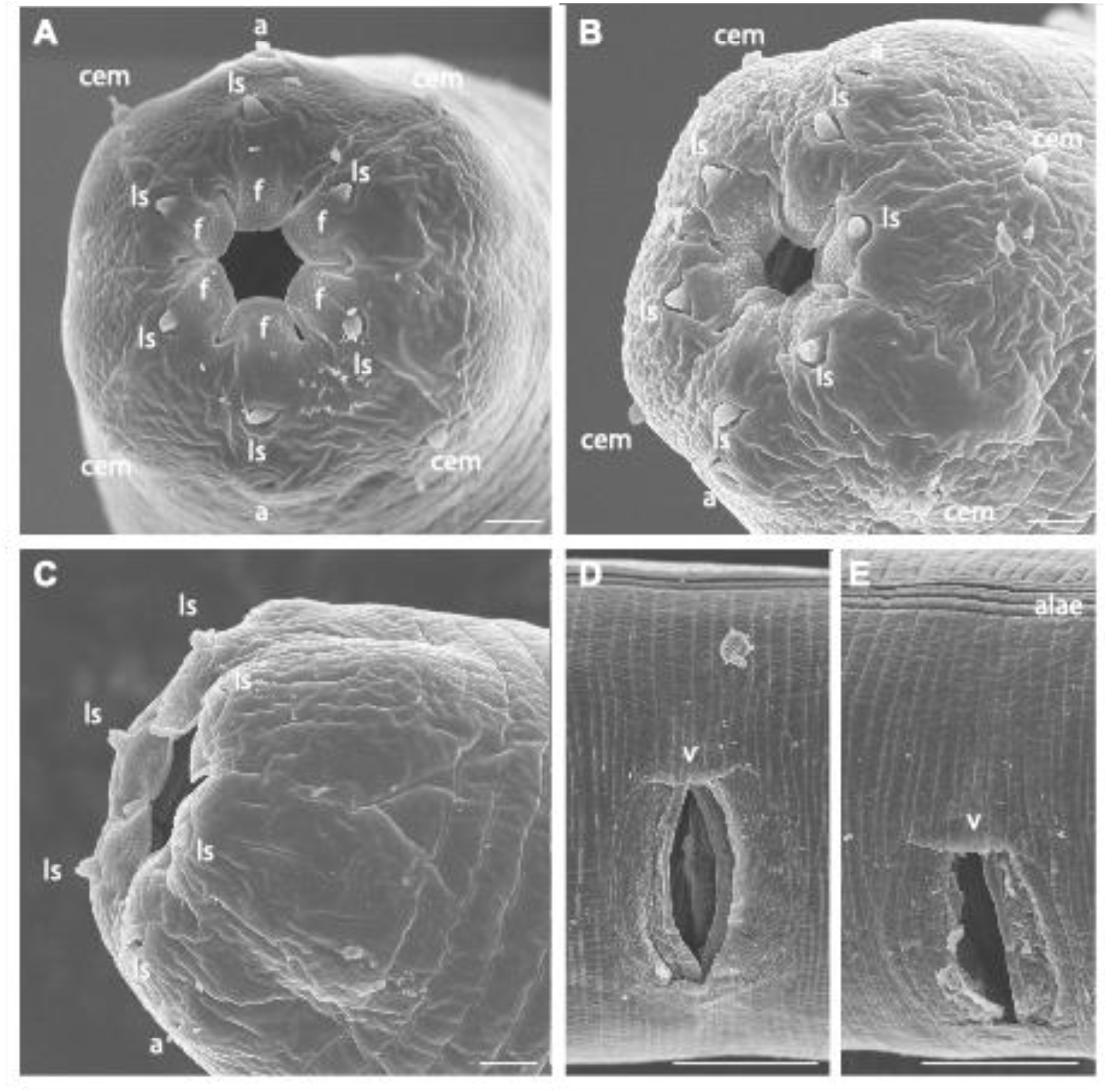
Scanning electron microscopy of heads and female vulva. (**A**): Head of adult male in anterior view. (**B**): Head of adult male in antero-lateral view. (**C**) Head of adult female in antero-lateral view (**D,E**): Female vulval openings in ventral views. In (**E**), alae are visible on the top of the micrograph. a: amphid; ls: labial sensillum; cem: male cephalic sensillum; f: flap on lips; v: vulva. Bars: (**A-C**) 1 µm; (**D,E**) 10 µm.

**Appendix Figure A2.**
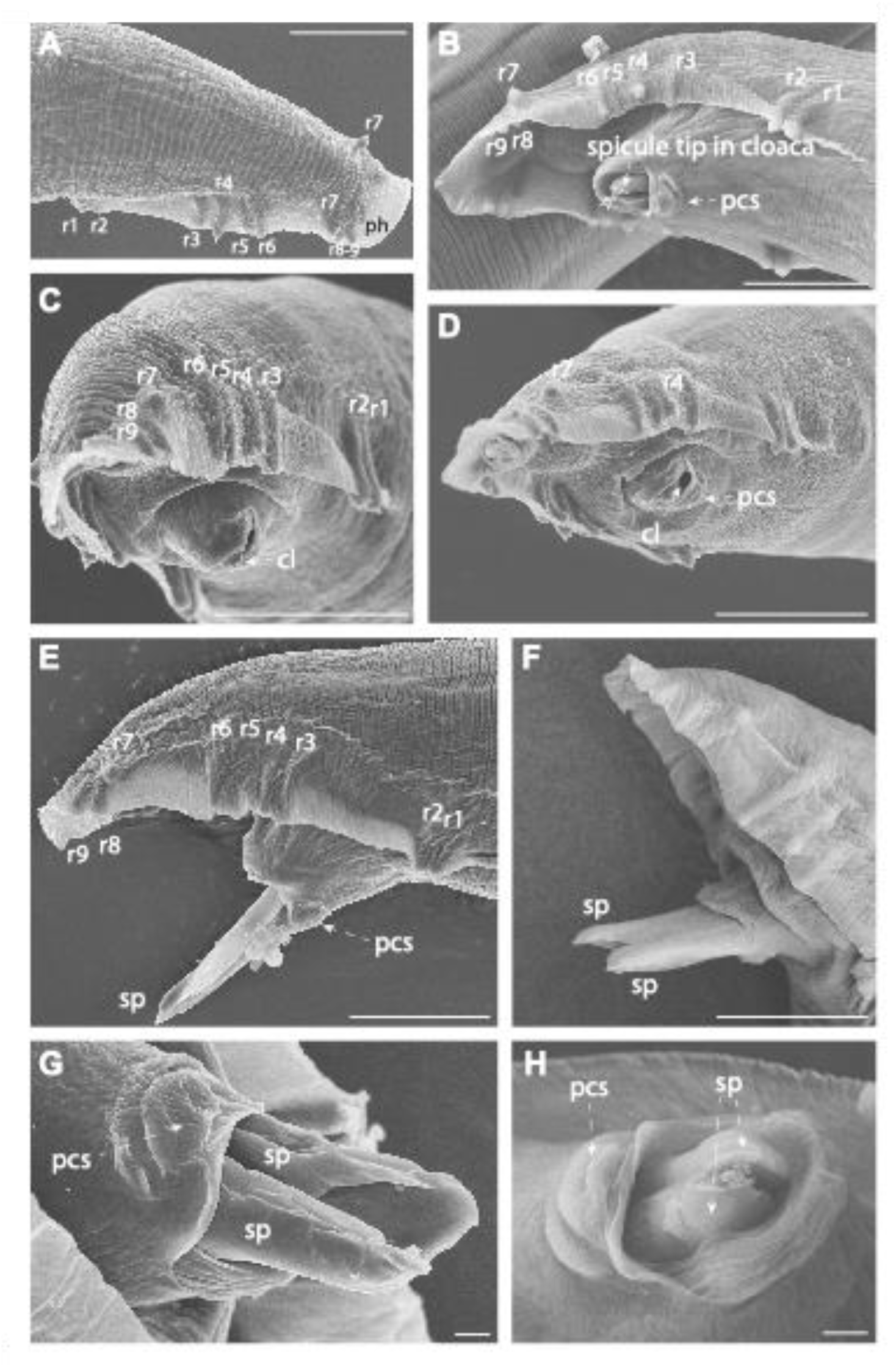
Scanning electron microscopy of male tails. (**A**) Dorsal view. (**B-D**) Ventro-lateral views, also from the posterior direction in (**C,D**). (**E,F**) Lateral views with extruded spicules. (**G,H**) Ventro-lateral views of the cloacal opening. Anterior is to the left in (**A,G,H**); to the right in (**B-F**). The male rays are numbered r1 to r9 according to antero-posterior position. The anterior dorsal ray is r4, and the posterior dorsal ray is r7. ph: phasmid. sp: spicule: pcs: precloacal sensillum. Bars: (**A-F**) 10 µm; (**G-H**) 1 µm.

## Notes

https://github.com/SerenaDingLab/Greenway_et_al_Caen-Invert-Intxns

### Competing Interest Statement

The authors have declared no competing interest.

### Summary of Updates

We have added more detail to the methods regarding the identification of the beetle species and nematode taxa, as well as the culturing conditions of the nematodes and methods for comparing the global distribution of species. We have added a description for C. sp. 8 as an Appendix, providing the species with the name C. apta to aid in clarity. Figure order revised.

